# Broad-spectrum antiviral activity of naproxen: from Influenza A to SARS-CoV-2 Coronavirus

**DOI:** 10.1101/2020.04.30.069922

**Authors:** Olivier Terrier, Sébastien Dilly, Andrés Pizzorno, Julien Henri, Francis Berenbaum, Bruno Lina, Bruno Fève, Frédéric Adnet, Michèle Sabbah, Manuel Rosa-Calatrava, Vincent Maréchal, Anny Slama Schwok

## Abstract

There is an urgent need for specific antiviral drugs directed against SARS-CoV-2 both to prevent the most severe forms of COVID-19 and to reduce viral excretion and subsequent virus dissemination; in the present pandemic context, drug repurposing is a priority. Targeting the nucleoprotein N of the SARS-CoV-2 coronavirus in order to inhibit its association with viral RNA could be a strategy to impeding viral replication and possibly other essential functions associated with viral N. The antiviral properties of naproxen, belonging to the NSAID family, previously demonstrated against Influenza A virus, were evaluated against SARS-CoV-2. Naproxen binding to the nucleoprotein of SARS-CoV2 was shown by molecular modeling. In VeroE6 cells and reconstituted human primary respiratory epithelium models of SARS-CoV-2 infection, naproxen inhibited viral replication and protected the bronchial epithelia against SARS-CoV-2 induced-damage. The benefit of naproxen addition to the standard of care is tested in an on-going clinical study.

---

The current pandemic of novel coronavirus disease 2019 (COVID-19), caused by severe acute respiratory syndrome coronavirus 2 (SARS-CoV-2) began in Wuhan, Hubei province, China, in December 2019 ^1,2^. As of April 22, 2020, there have been more than 2 475 723 confirmed cases of COVID-19, including 169 151 deaths in 213 countries, reported to WHO (https://www.who.int/emergencies/diseases/novel-coronavirus-2019). SARS-CoV-2 is a beta-coronavirus closely related to the severe acute respiratory syndrome coronavirus-1 (SARS-CoV-1) and the Middle East respiratory syndrome coronavirus (MERS-CoV), that emerged in 2003 and 2012, respectively. These viruses are also transmitted from animals to humans and cause severe respiratory diseases in afflicted individuals.

There is an urgent need for specific and effective antiviral drugs directed against pandemic SARS-CoV-2 to prevent the most severe forms of COVID-19. In that context, drug repurposing is a priority. Ongoing advances in our knowledge of this new virus and its pathogenesis are revealing an exacerbated inflammatory response in severe COVID-19 cases, with a similar cytokine storm observed in severe cases of H5N1 Influenza virus infection and 1918 Influenza A pandemics^3^. In that regard, the symptoms of respiratory distress caused by COVID-19 could be reduced by drugs combining anti-inflammatory and antiviral effects^4^.

Several current therapeutic approaches involve drug repositioning, in light of what has been achieved for other viruses, particularly influenza viruses^5^. Our previous work showed that naproxen, an approved anti-inflammatory drug, is an inhibitor of both cyclooxygenase (COX) and of Influenza A virus nucleoprotein ^6^. Although NSAIDs were the subject of a precautionary measure by the French Ministry of Health in mid-March 2020 regarding the use of these drugs in the event of COVID-19 infection, there is currently no evidence of any particular toxicity of this family of drugs ^7-8^. Moreover, the ability of naproxen to decrease viral load and inflammation through the inhibition of cyclooxygenase activity could be beneficial for limiting the cytokine storm that may happen in severe forms of COVID19. Importantly, naproxen has the advantage of being a generic drug, readily available and often used for other indications than COVID-19 by fragile populations.

Naproxen binding to the Influenza A virus nucleoprotein blocked viral RNA association with the nucleoprotein and impeded its self-association ^6, 9^; consequently, naproxen strongly reduced viral transcription/ replication in infected cells and protected mice against an infection with Influenza A virus ^6, 10^. Viral nucleoproteins are unique to the virus and no equivalent of these proteins are found in the host cell, which makes them attractive targets for potential antivirals ^12-13^. Moreover, the viral nucleoprotein is an important diagnostic marker of infection with SARS-COV and /or Influenza A ^14-15^. SARS-CoV-1 virions contains multiple copies of the N protein (ca 1000 copies) located inside the particle, thus N is one of the most abundant structural proteins in CoVs ^16^. Through its interaction with the coronavirus membrane (M) protein, the N protein drives virus assembly and budding. N binds to the long viral RNA genome to form a virion core comprising a ribonucleoprotein (RNP) complex that assumes a long helical structure. The RNP is important for replication and transcription in which the N protein also plays a critical role. Interactions between N and the non-structural protein-3 are also important for replication. In addition to its role in different aspects of the viral cycle, the N protein not only highjacks cellular processes, including the progression of the host cell cycle and apoptosis, but also modulates the immune response, by for example inhibiting the interferon response^17^.

In this study, we combined structure-based modeling of naproxen binding to the nucleoprotein of SARS-CoV-2 with experimental approaches to explore and validate whether naproxen harbors antiviral activity against SARS-CoV-2 pandemic virus, as previously implemented for Influenza A virus^9-10, 18^.

## Structure-based modeling of naproxen binding to the nucleoprotein of SARS-CoV-2

The nucleoproteins N of enveloped, positive-sense, single-stranded viruses Coronavirus (CoV) share with negative-sense single-stranded viruses such as Influenza A virus the ability to bind to- and protect genomic viral RNA without sequence specificity and to form self-associated oligomers ^12-13, 18-19^. Despite the limited sequence similarity between them, the N-terminal domains of three coronaviruses SARS-Cov-1, MERS-CoV, SARS-CoV-2 and Influenza A virus all presented a wide, positively-charged groove in which the viral negatively-charged RNA binds, Figure 1 (A-D)^20-21,13^. The coronavirus N proteins present a central antiparallel sheet of 3-5 β-strands structure and flexible linkers and loops, with emphasis on the large extrusion carrying many basic residues, which likely helps adjust binding to the RNA. Such flexibility ^22^ was evaluated here from the crystallographic b factor (Figure 1 E-H). The central parts of the β -sheet of these CoV proteins concentrate most of the aromatic residues (Figure 1I). For instance, SARS-CoV-2 nucleoprotein amino-terminal domain counts 14 aromatic residues among a total of 124 modeled in 6VYO.pdb. The consecutive, 5-aromatics pentapeptide ^108^WYFYY^112^ is located at the center of the electropositive groove (Figure 1I, red arrow). This contrast with the nucleoprotein of Influenza A virus which has a single aromatic residue (Y148) within its RNA binding groove in which naproxen (Figure 1J) was shown to bind, making electrostatic (blue arrow) and hydrophobic interactions (red arrow) with conserved residues of the RNA binding groove and C-terminal domain (Sup. Figure 1)^6^. From these observations, we could expect that the N-terminal domain of CoVs N may offer more than a single binding site for naproxen, which could concentrate within the cluster of aromatic residues found in the middle of the β-sheet. This is exactly what is observed in SARS-CoV-1 (Figure 1K, N and supplementary Figure 1C-K). The recently published structure of MERS-CoV N terminal domain and the SARS-Cov-2 N-terminal domain were crystallized as a dimer of dimer, which offers a potential site for naproxen at the interface of the dimer ^13^ in addition to a binding site on one monomer of the protein. These two sites are shown in Sup. Figure 1E, F. Naproxen stabilized this “non -natural” interface as found in the crystal structure for the designed ligands by Hou et al^13^. Figures 1L, O and M, P and Sup. Figure 1H, E indeed show that naproxen can also bind at this interface in both MERS-CoV and SARS-Cov-2, respectively, but an additional binding site in SARS-CoV-2 monomer was also identified (Sup. Figure 1F). Supplementary Figure 1A-D shows that naproxen binds at the interface between hydrophobic and hydrophilic parts of N proteins, to account for the hydrophobic interactions of its aromatic core with usually aromatic residues (π - stacking) and methyl group of the methoxy group with hydrophobic apolar residues and polar and/or electrostatic interactions via its carboxylate moiety. Accordingly, the following order of free binding energies of naproxen binding to the N-terminal domain of N were calculated: SARS-CoV-1 (ΔG= -30 Kcal/mol) ≈MERS-CoV (ΔG= -29 Kcal/mol) ≈ SARS-Cov-2 (ΔG= -26 to - 32) Kcal/mol for site 1 and 2, respectively. This suggests a similar affinity of naproxen for SARS-CoV-1 and MERS-CoV, similar or slightly lower affinities of naproxen for SARS-CoV-2 than naproxen affinity for Influenza A virus based on a free energy calculation of (ΔG= -30 Kcal/mol). Taking together, this work suggests naproxen binding in the N-terminal domains of SARS-CoV-2 N protein and likely to the SARS-CoV-1 and MERS-CoV N in similar binding modes. It is likely that this binding would impede viral RNA binding to N as naproxen binding sites on the N-terminal domain of N are found in the RNA binding groove. Competing with N binding to RNA would repress replication and decrease the level of RNA produced by the virus. The experimental quantification of the viral RNA by **RT-qPCR** tests this hypothesis in the below antiviral assays.

**Figure 1:**
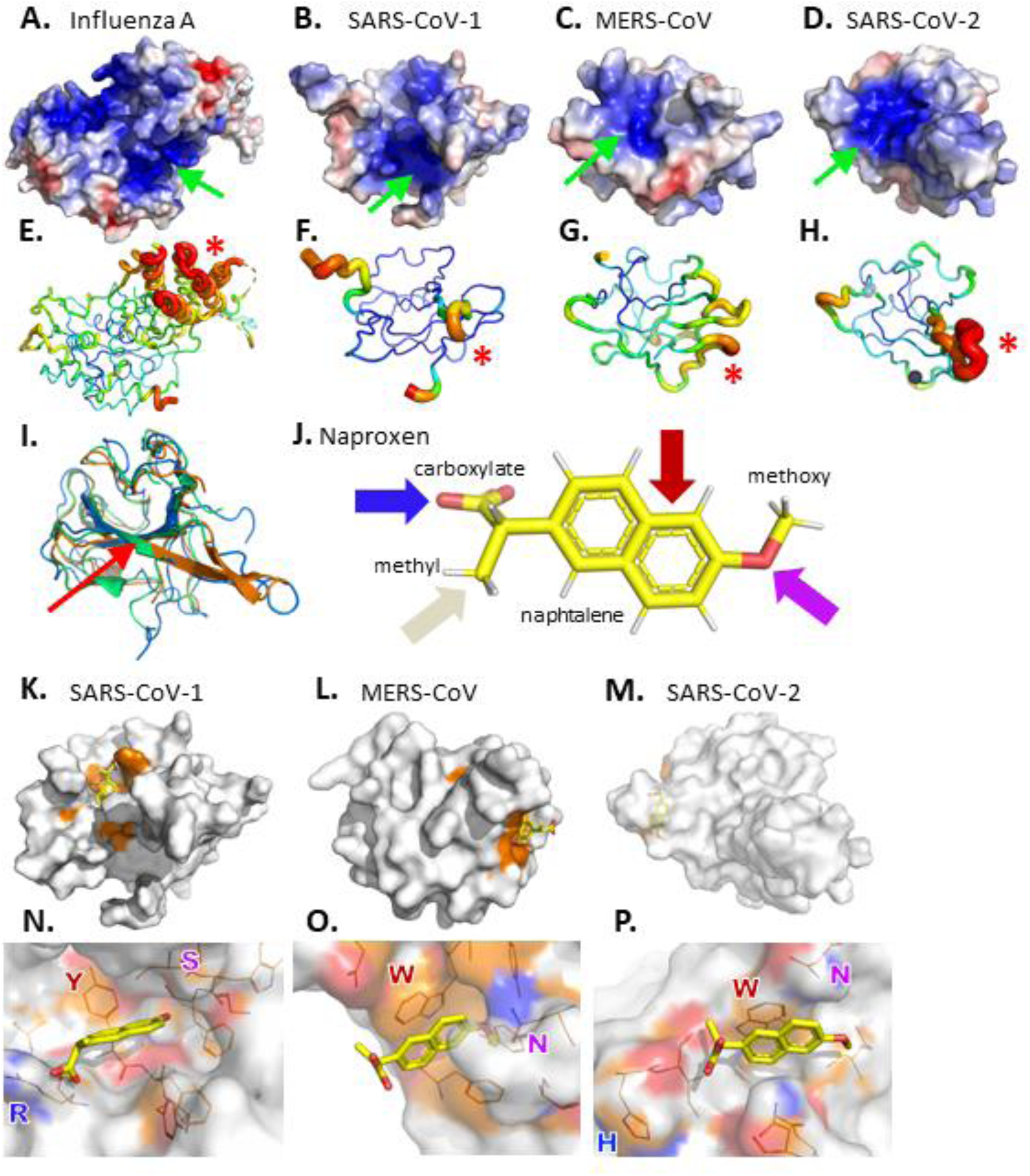
Structural comparison of viral nucleoproteins N of (A.) influenza A 2IQH.pdb and of amino-terminal domains of (B.) SARS-CoV-1 1SSK.pdb (C.) MERS-CoV 6KL2.pdb (D.) SARS-CoV-2 6VYO.pdb. Electrostatic surface potentials are computed according to Adaptive Poisson-Boltzmann Solver Electrostatics plugin from PyMOL in standard parameters. Electronegative potentials are colored red while electropositive potentials are colored blue. A large electropositive cleft is visible on each protein, pointed with a green arrow; it maps the putative RNA interaction sites. (E.F.G.H.) Main chains flexibilities are approximated as crystallographic b-factors, with main chain ribbon diameter proportional to b-factor value; red is highest b-factor while blue is lowest b-factor. Most mobile elements are identified by a red asterisk (*); they group on one external side of the protein, facing solvent. Mobile elements of highest b-factors are proximal to electropositive surfaces, supporting an interaction scenario where (1) RNA docks onto an electro-complementary surface before (2) the protein conformation is induced into the N-RNA complex of highest stability. (I.) Pairwise structural alignment of equivalent Carbons α of SARS-CoV-1 (blue), MERS-CoV (green) and SARS-CoV-2 (orange) nucleoproteins. The red arrow emphasizes the conserved RWYFYY sequence (J.) Naproxen tridimensional structure in sticks. Frequent residue types in bonding distances in all docking poses are highlighted with arrows: blue, charged electrostatic; red, aromatic; purple, uncharged polar; teal, van der Waals contact. (K.L.M.) Naproxen modelled binding site on amino-terminal RNA interacting domains of (K.) SARS-CoV-1 SSK1.pdb, (L.) MERS-CoV 6KL5.pdb, (M.) SARS-CoV-2 6VYO.pdb. Connoly solvent excluded surfaces are colored white. Residues at a 4 Å distance to naproxen are colored orange. (N.O.P.) Close-up view of docking pockets with residues at a 4 Å distance to naproxen represented in lines. Nitrogen is colored blue, oxygen is colored red. Images were ray traced with PyMOL version 2.0.6. Orientations were selected to highlight discussed properties.

## Naproxen inhibits SARS-CoV-2 infection in VeroE6 cells and in reconstituted human airway epithelia (HAE)

Based on the docking analysis described above, we then evaluated the potential anti SARS-CoV-2 activity of naproxen *in vitro*. As shown in **Figure 2A-B**, naproxen efficiently reduces viral replication in African green monkey VeroE6 cells in a dose-response manner. A unique treatment at 1hour post-infection (hpi) resulted in 50% inhibitory concentration (IC50) values of 46.07 µM at 48 hpi and 59.53 µM at 72 hpi. We determined a cytotoxic concentration CC50 > 1000 µM. This value compares well with the CC50 previously determined in MDCK and A549 cells to be 1605 ± 300 µM^6^. The selectivity index (SI) was SI= CC50/IC50, SI > 21.71 and 19.79, respectively.

**Figure 2:**
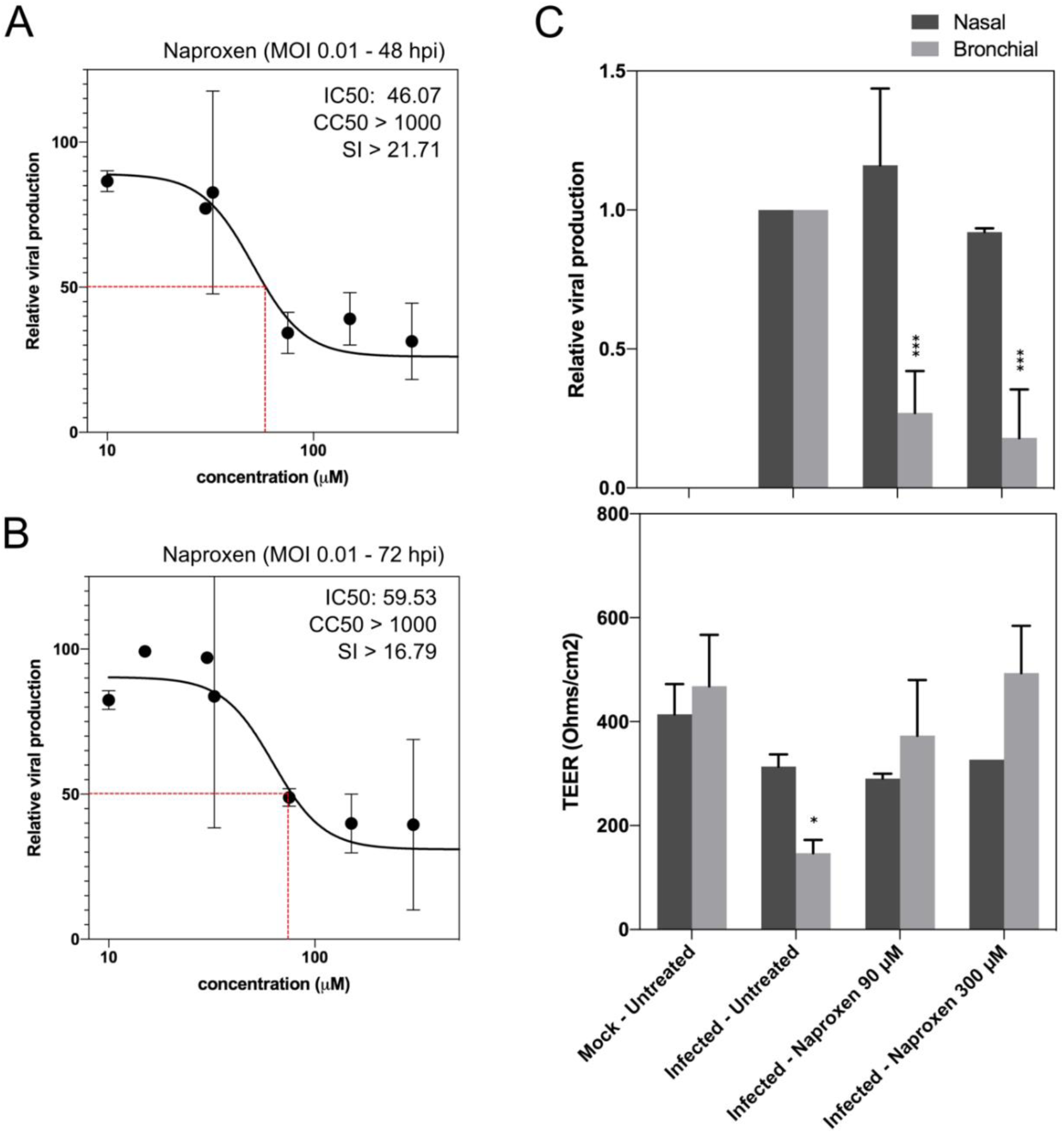
Naproxen inhibits SARS-CoV-2 infection in VeroE6 cells and in HAE. (A-B) Dose-response curves of naproxen at 48 and 72 hpi in infected VeroE6 cells. (C) Relative intracellular viral genome quantification and trans-epithelial resistance (TEER in Ohms/cm2) between the apical and basal poles in nasal and bronchial HAE at 48 hpi. Results are expressed in relative viral production compared to the infected untreated control and relative TEER compared to t=0 (before infection). ***P <0.001 and *P <0.05 compared to the infected untreated (viral titers) or uninfected (TEER) groups by Student t-test. Data are representative of three independent experiments.

We further evaluated the antiviral effect of naproxen using a more biologically relevant experimental model, notably the nasal and bronchial MucilAir™ reconstituted human airway epithelia (HAE)^23^. Developed from biopsies of nasal or bronchial cells differentiated in the air/liquid interphase, these models reproduce with high fidelity most of the main structural, functional and innate immune features of the human respiratory epithelium that play a central role in the early stages of infection and constitute robust surrogates to study airway disease mechanisms and for drug discovery^23^. Post-infection treatment of nasal HAE with 90 or 300 µM naproxen did not show an antiviral effect at 48 hpi compared to the mock-treated control (**Fig. 2C**, upper panel).

Conversely, significant reductions in intracellular SARS-CoV-2 viral titers were observed for the two treatment conditions in bronchial HAE (73 and 82% reduction vs mock-treated control, respectively). This reduction in viral titers correlated with naproxen inducing a protective effect of the bronchial epithelium integrity, as shown by Trans-epithelial electrical resistance (TEER), considered as a surrogate of epithelium integrity. TEER values in both naproxen groups were comparable to those of the uninfected and untreated control (**Fig. 2C**, lower panel) and significantly higher than those of the untreated control (Fig. 2C, lower panel). The antiviral effects of naproxen in reconstituted human bronchial epithelium are consistent with the IC50 value determined in Vero E6 cells. However, the lack of antiviral effect of naproxen on nasal epithelium is puzzling. The viral load is usually lower in the nasal cavity as compared to the lungs^16, 24^. The nasal epithelium is an important portal for initial infection, and may serve as a key reservoir for viral spread across the respiratory mucosa and an important locus mediating viral transmission. The number of host cells as pneumocytes and alveolar macrophages permissive to viral entry is higher in the lungs than the mucous cells in the nasal cavity. Moreover, recent interactome analysis support the hypothesis that SARS-CoV-2 preferentially hijacks host proteins available in lung tissue, and N targets stress granule protein G3BP1, an essential antiviral protein which is known to induce innate immune response ^7^.

Interestingly, we previously determined that the peak of viral replication was reached earlier in bronchial (48-72 hpi) than in nasal HAE, in which a progressive increase in infectious viral titers was observed until at least 96 hpi^23^. Using this model, this differential antiviral effect between airway sites was observed for naproxen in this work and for remdesivir in a recent report that indicates an antiviral effect mainly observed in the lower respiratory tract of non-Human primates^23^. It is interesting to note that naproxen is found very protective against viral-induced damages at 48 h post-infection in the reconstituted bronchial epithelium (Figure 2C and Figure 4 in ^23^). Therefore, we speculate that the lack of naproxen effect in nasal epithelium as opposed to its antiviral effect in pulmonary epithelium could be associated with differential expression or activity of host factor(s), and/or with slower replication kinetics in the nasal vs pulmonary epithelia. FAD-approved drugs including nitazoxanide, chloroquine and remdesivir exhibited IC50s in the low micromolar range against a clinical isolate of COVID-2019 in vitro^25^. Their corresponding selectivity index were 17, 129 and 89, respectively and CC50 values found to be 36 µM and in the > 100 µM range. Here, the selectivity index determined for naproxen for SARS-CoV-2 in VeroE6 cells is SI > (19 ± 3) µM.

The IC50 of naproxen for SARS-CoV2 compares well to IC50 = (25 ± 7) µM and SI = 64 of naproxen determined for Influenza A at the same MOI = 10^−2^. These values are consistent with the determination of the free energies of naproxen binding to N. We cannot exclude additional putative naproxen binding sites of naproxen in the C-terminal domain of SARS-CoV2 or in the unstructured serine-rich domain that could enable naproxen to further interfere with N functions. In addition to the antiviral activity against SARS-CoV-2 virus demonstrated here, naproxen also exhibited antiviral properties in cellular and rodent models of Influenza A and B ^6, 10, 26^. Naproxen also inhibited the replication of the unrelated Zika virus ^27^. Therefore, from three unrelated single-stranded RNA virus, we suggest that naproxen could present broad-spectrum antiviral properties. Moreover, the combination of clarithromycin, naproxen and oseltamivir reduced mortality of patients hospitalized for H3N2 Influenza infection when compared to oseltamivir alone in a phase IIB/III clinical trial ^11^.

## Conclusion

We suggest in this study that one of the NSAIDs, naproxen, has a direct antiviral activity on SARS-CoV2 by its effect on the nucleoprotein-mediated replication and possibly via other functions of N, as inhibition of interferon^17^. Indeed, the cellular assays demonstrated subsequent inhibition of viral replication by naproxen, with recovery of intact pulmonary epithelium protected from the pandemic virus. These very encouraging results prompted us to test in a clinical trial, named ENACOVID, the effect of addition of naproxen on the standard of care in severely-ill patients. This trial is supported by APHP, Assistance Publique, Hôpitaux de Paris areas and deposited on March 26, https://www.clinicaltrialsregister.eu/ctr-search/search?query=eudract_number:2020-001301-23.

## Experimental

### Modeling

The following X-ray structures extracted from the Protein Data Bank (PDB) have been used: N-terminal domain of SARS-CoV-2: 6VYO ^20^, SARS-CoV-1: 1SSK ^21^, MERS-CoV: 6KL2/ 6KL5 ^13^; The Influenza A nucleoprotein PDB 2IQH was used as a comparison^28^. The binding protocol and subsequent MD are described in the Supplementary data.

### Material

Naproxen was purchased from Sigma (reference N5160)

### Virus

All experiments involving the manipulation of infectious SARS-CoV-2 were performed in biosafety level 3 (BSL-3) facilities, using appropriate protocols. The BetaCoV/France/IDF0571/2020 SARS-CoV-2 strain used in this study was isolated directly from a patient sample as described elsewhere^23^. (https://www.biorxiv.org/content/10.1101/2020.03.31.017889v1). The dose-response antiviral evaluation of naproxen in Vero E6 cells and in reconstituted human airway epithelia (HAE) were performed as described before https://www.biorxiv.org/content/10.1101/2020.03.31.017889v1 and further detailed in the Supplementary data.

## Funding

This Consortium is grateful to Sorbonne University for granting our project CovNucleovir on a Flash COVID funding of the Medicine Faculty. Professor Frederic Adnet is gratefully acknowledging Assistance Publique-Hôpitaux de Paris APHP for granting the ENACOVID clinical trial. Part of this work was funded by INSERM REACTing (REsearch & ACtion emergING infectious diseases), CNRS, and Mérieux research grants (TO and MRC, VirPath team). We are grateful to Drs HO Bertrand and K. Luckman from Biovia for the gift a temporary licence for Discovery Studio (Dassault Systemes) to SD.

## Supplementary methods

### Modeling

The binding mode of naproxen for the protein structures was assessed in a two-step protocol. First, the binding site was defined by blind docking to the whole protein using the docking program AutoDock Vina ^34^. The docking space was defined visually in order to encompass the proteic model with a 2 Å margin at least as a tetragonal box. Three replicas with the default scoring function Vina^29^ were performed to check the convergence of the experiment. The relevance of the binding sites identified by blind docking was confirmed by the cavity detection algorithm of Discovery Studio version 2020, which detected similar sites. A site-docking was then performed using Libdock with energy minimization using smart minimizer (Discovery Studio version 2020) to more precisely identify the naproxen binding mode. In this docking, three replicas were also applied but on a restricted space, encompassing the cavity where the ligand was blind docked. For each protein, the most representative pose of naproxen (i.e. the pose more often found) was selected. The resulting protein-NP complexes were finally refined by a molecular dynamics simulation using the CHARMm force field ^35^ and the standard dynamic cascade protocol of Discovery studio version 2020. The free energies of naproxen binding to the various structures were evaluated using implicit Distance-dependent dielectrics solvent model found in Discovery Studio version 2020.

### Dose-response antiviral evaluation in Vero E6 cells

Multi-well 6 plates were seeded with VeroE6 cells 24 h before infection (day -1). On day 0, 90%-confluent wells were washed twice with PBS and infected with SARS-CoV-2 at a multiplicity of infection (MOI) of 0.01. One hour after infection, the inoculum was removed and cells were subsequently treated with the indicated dilutions of naproxen in DMEM. DMEM was used as mock-treatment control. Plates were then incubated at 37 °C and 5% CO2. Supernatants were collected at 48 and 72 hpi and stored at -80 °C for viral RNA extraction with the QiAmp viral RNA Kit (Qiagen) and titration by RT-qPCR as described elsewhere^21^. Drug 50% cytotoxic concentration (CC50), 50% inhibitory concentration (IC50) and selectivity index (SI) values were calculated using the Quest Graph IC50 calculator (AAT Bioquest).

### Evaluation of antiviral activity in reconstituted human airway epithelia (HAE)

MucilAir™ reconstituted HAE were obtained from Epithelix SARL (Geneva, Switzerland) and maintained in air-liquid interphase with specific culture medium in Costar Transwell inserts (Corning, NY, USA) according to the manufacturer’s instructions. After two washes with warm OptiMEM medium (Gibco, ThermoFisher Scientific), apical poles were inoculated with a 150 μl dilution of virus in OptiMEM medium at a MOI of 0.1. OptiMEM was used as mock-infection control. One hour after viral infection, MucilAir™ culture medium containing or not (untreated) specific dilutions of naproxen in were applied through the basolateral poles. Treatments were repeated at 24 hpi. At 48 hpi, HAE cells were harvested in RLT buffer (Qiagen) and total ARN was extracted using the RNeasy Mini Kit (Qiagen) and stored at -80 °C for subsequent titration by RT-qPCR. Variations in transepithelial electrical resistance (Δ TEER) were measured using a dedicated volt-ohm meter (EVOM2, Epithelial Volt/Ohm Meter for TEER) and expressed as Ohm/cm^2^.

**Supplementary Figure 1:**
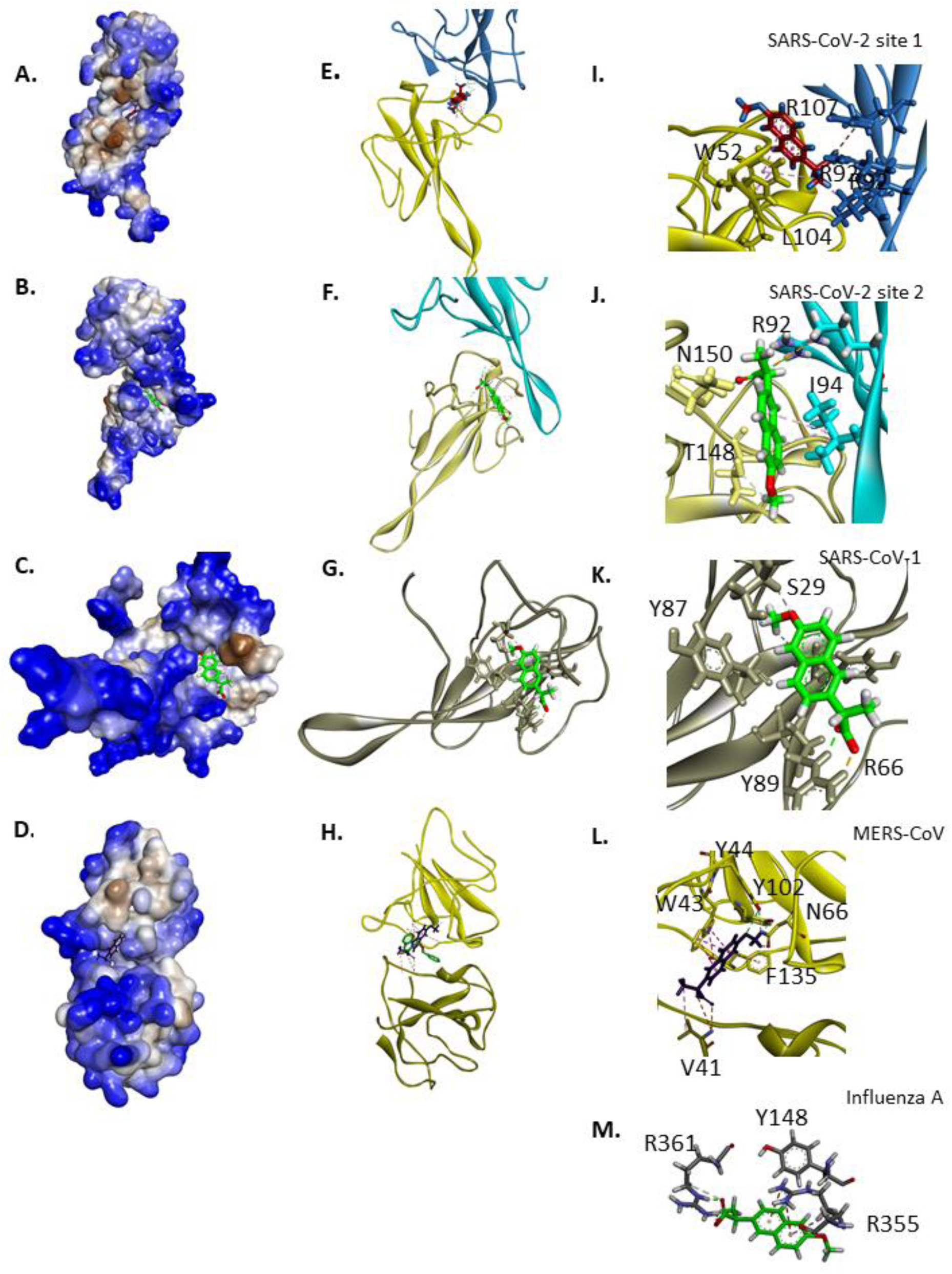
Each structure is represented in hydrophobic (brown) to hydrophilic (blue) surface (first column, in ribbon (second column), a close-up is shown in the third column. **Binding of naproxen to the N-terminal domain of SARS-CoV2 site 1** (most frequent site): panels A, E and I: the carboxylate interacted by polar interactions with S105 and electrostatic interactions with R92, R107, the latter belonging to the sequence R^107^WYFYY^112^ of monomer D, L104 of monomer D made hydrophobic contacts with the methyl of the methoxy group, the aromatic core of naproxen made hydrophobic contacts with Y112 (of the sequence R^107^WYFYY^112^) and stacked on W52. **Binding of naproxen to the N-terminal domain of SARS-CoV2 site 2:** panels B, F and J: the carboxylate interacted by polar and electrostatic interactions with N150 (monomer A) and R92 (monomer D), respectively, T148 of monomer A made hydrophobic contacts with the methyl of the methoxy group, the aromatic core of naproxen made hydrophobic contacts with I94 (monomer D). **Binding of naproxen to the N-terminal domain of SARS-CoV1:** panels C, G and K: the carboxylate interacted by electrostatic interactions with R66, the oxygen of the methoxy group made polar interaction with S29, the aromatic core of naproxen was in hydrophobic/ stacking interactions with surrounding Y87 and Y89 of the sequence R^85^WYFYY^90^. **Binding of naproxen to the N-terminal domain of MERS-CoV:** panels D,H and L: the carboxylate interacted by polar interactions with N66 (monomer D), the methyl group formed hydrophobic contacts with V41 (monomer A), the aromatic core of naproxen stacked on W43 and F135 (monomer D), Y44 and Y102 of the sequence R^97^WYFYY^102^ (monomer D) made both hydrophobic interactions and polar interactions (via its OH group) with the methoxy group of naproxen. The ligand crystallized in this structure is depicted in green for comparison. Panel M shows naproxen binding to Influenza A N that involved electrostatic interactions of the carboxylate with R361, a cation -π interaction with R355 and hydrophobic interaction with Y148 and interactions of the last C-terminal F (not shown) with the methyl of the methoxy group.

## Notes

### Competing Interest Statement

The authors have declared no competing interest.

